## Supplementary information for "Ligand-directed two-step labeling to quantify neuronal glutamate receptor trafficking"

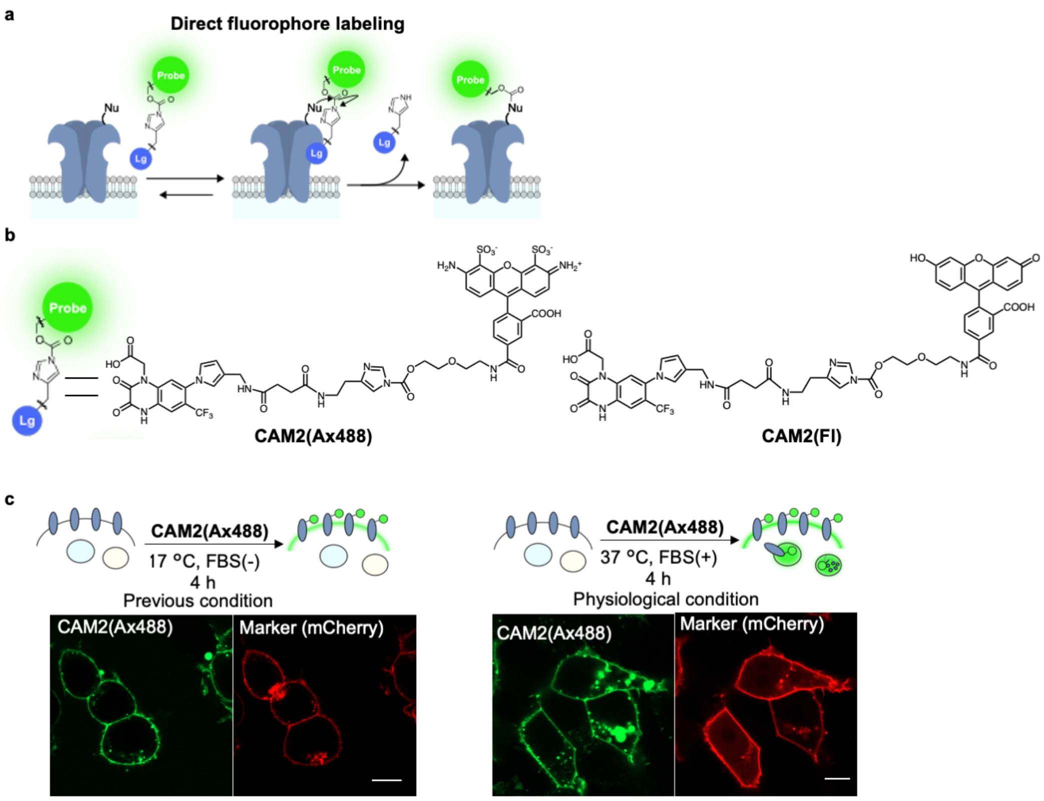


**Supplementary Figure 1 | Direct fluorophore labeling to AMPARs using CAM2(Ax488) or CAM2(Fl) in HEK293T cells.** (**a**) Schematic illustration of direct fluorophore labeling to AMPARs using CAM2 reagent. Lg, selective ligand for AMPARs; Nu, nucleophilic amino acid residue. (**b**) Chemical structure of **CAM2(Ax488)** or **CAM2(Fl)**. (**c**) Confocal live imaging of the HEK293T cells labeled with 2 μM **CAM2(Ax488)** under previous condition (in left) (ref S1) or under physiological cell culture condition (in right). In left, chemical labeling was conducted in serum-free medium at 17 °C. In right, chemical labeling was conducted in growth medium containing 10% FBS at 37 °C. mCherry-F was utilized as a transfection marker. Scale bars, 10 μm.

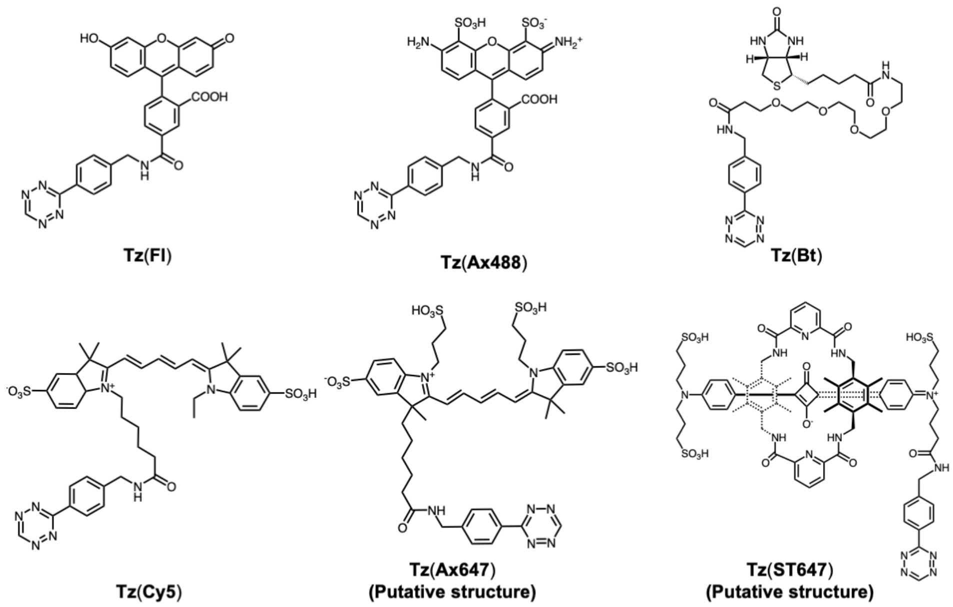


**Supplementary Figure 2 | Detailed chemical structure of Tz probes.**

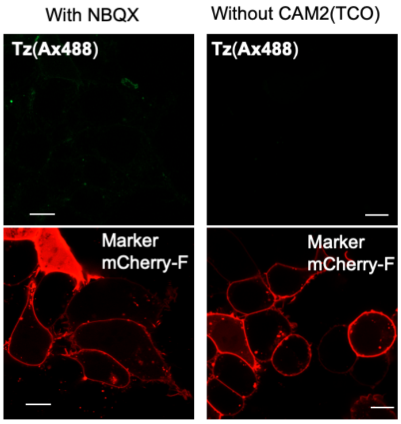


**Supplementary Figure 3 | Control experiment for the confocal live imaging of the HEK293T cells expressing AMPARs labeled with 2 µM CAM2(TCO) and 0.1 µM Tz(Ax488).** Co-presence of NBQX or absence of CAM2(TCO) hampers fluorescent labeling to cell-surface AMPARs in the two-step labeling. See also Figure 2b. mCherry-F was utilized as a transfection marker. Scale bars, 10 µm.

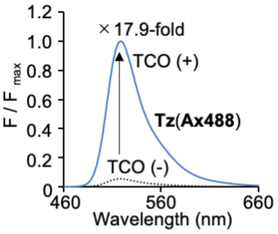


**Supplementary Figure 4 | Turn-on type fluorescent property of Tz(Ax488) after addition of TCO-PEG4-COOH.** Fluorescent spectra before and after addition of TCO-PEG4-COOH. [**Tz(Ax488)**] = 0.1 μM. [TCO-PEG4-COOH] = 1 μM. Excitation wave length is 430 nm.

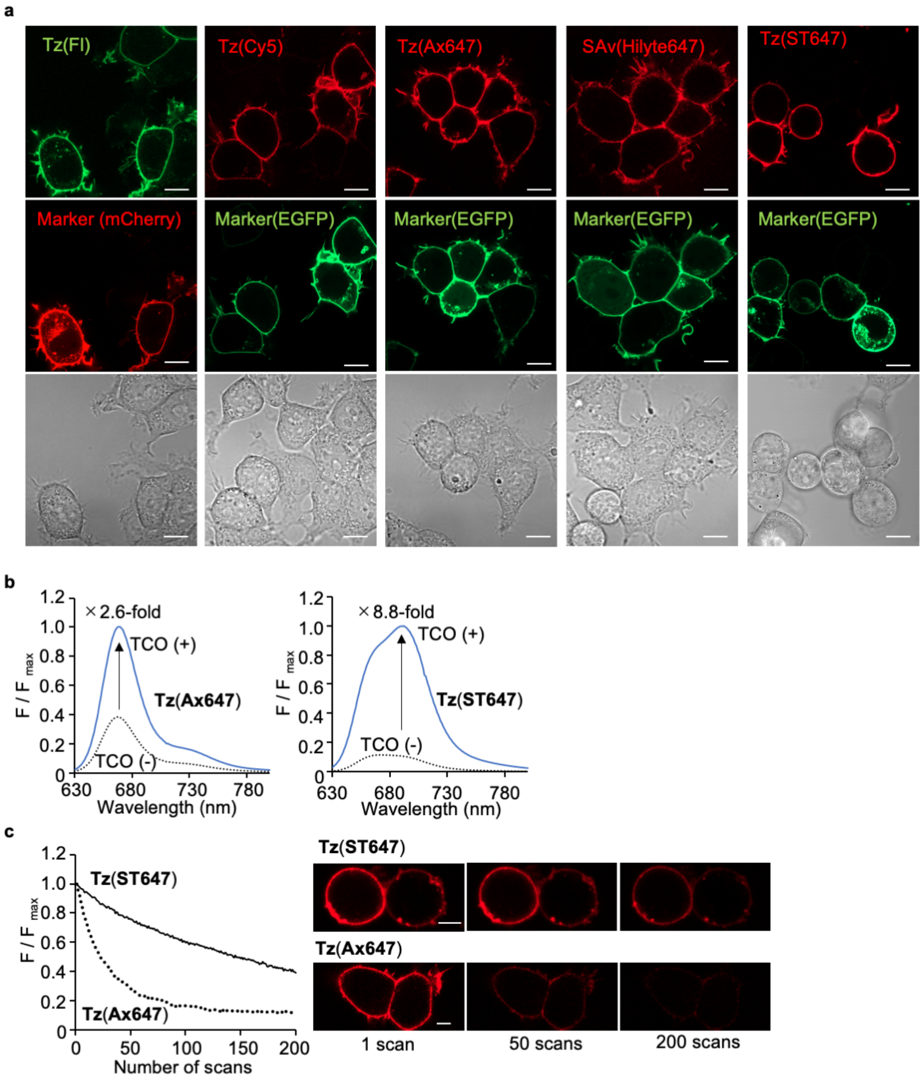


**Supplementary Figure 5 | Tethering various kinds of Tz-probes for visualization of cell-surface AMPARs in HEK293T cells** (**a**) Confocal live imaging of the HEK293T cells labeled with 2 µM **CAM2(TCO)** and 0.1 µM of each Tz-probe. Labeling was conducted as described in Figure 2b. In the case of the **Tz(Bt)** labeling, SAv(Hilyte647) was added for visualizing biotin-labeled AMPARs. mCherry-F or EGFP-F was utilized as a transfection marker. Scale bars, 10 µm. (**b**) Turn-on type fluorescent property of **Tz(Ax647)** or **Tz(ST647)** after addition of TCO-PEG4-COOH. Fluorescent spectra before and after addition of TCO-PEG4-COOH. [**Tz(Ax647)** or **Tz(ST647)**] = 0.1 µM. [TCO-PEG4-COOH] = 1 µM. e.x. = 610 nm. (**c**) Photostability of SeTau-647 labeled to cell-surface AMPARs in HEK293T cells. Photostability of SeTau-647 or Alexa 647 labeled to cell-surface AMPAR were evaluated by confocal live cell imaging. This result indicates that SeTau-647 has high photostability compared with Alexa 647. Scale bars, 5 µm.

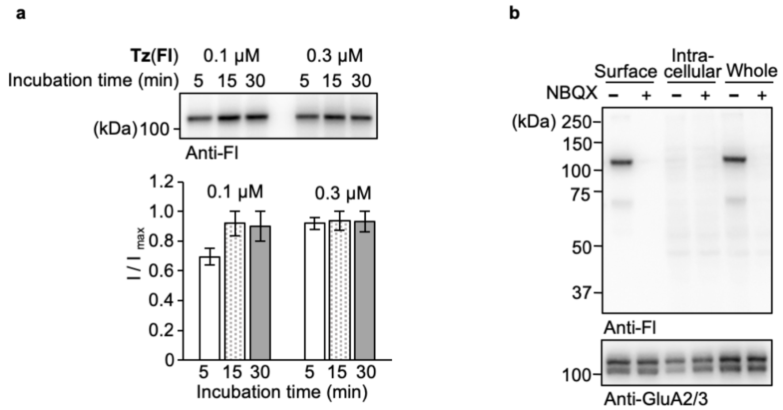


**Supplementary Figure 6 | Reaction kinetics of tetrazine ligation in cell lysates and a whole blot for surface, intracellular or whole-cell labeling in HEK293T cells.** (**a**) Reaction kinetics of tetrazine ligation in cell lysate evaluated by western blotting. After cell lysis of **CAM2(TCO)**-labeled HEK293T cells, each concentration of **Tz(Fl)** was added for 5, 15 or 30 min. Then, 10 μM TCO-PEG4-COOH was added for quenching **Tz(Fl)**. This result indicates that the tetrazine ligation was saturated within 15 min. (**b**) Whole blot for surface, intracellular or whole-cell. The sample was prepared as described in Figure 3g. Selective band corresponding to AMPAR was observed, which indicates high selectivity of the tetrazine ligation even in cell lysates.

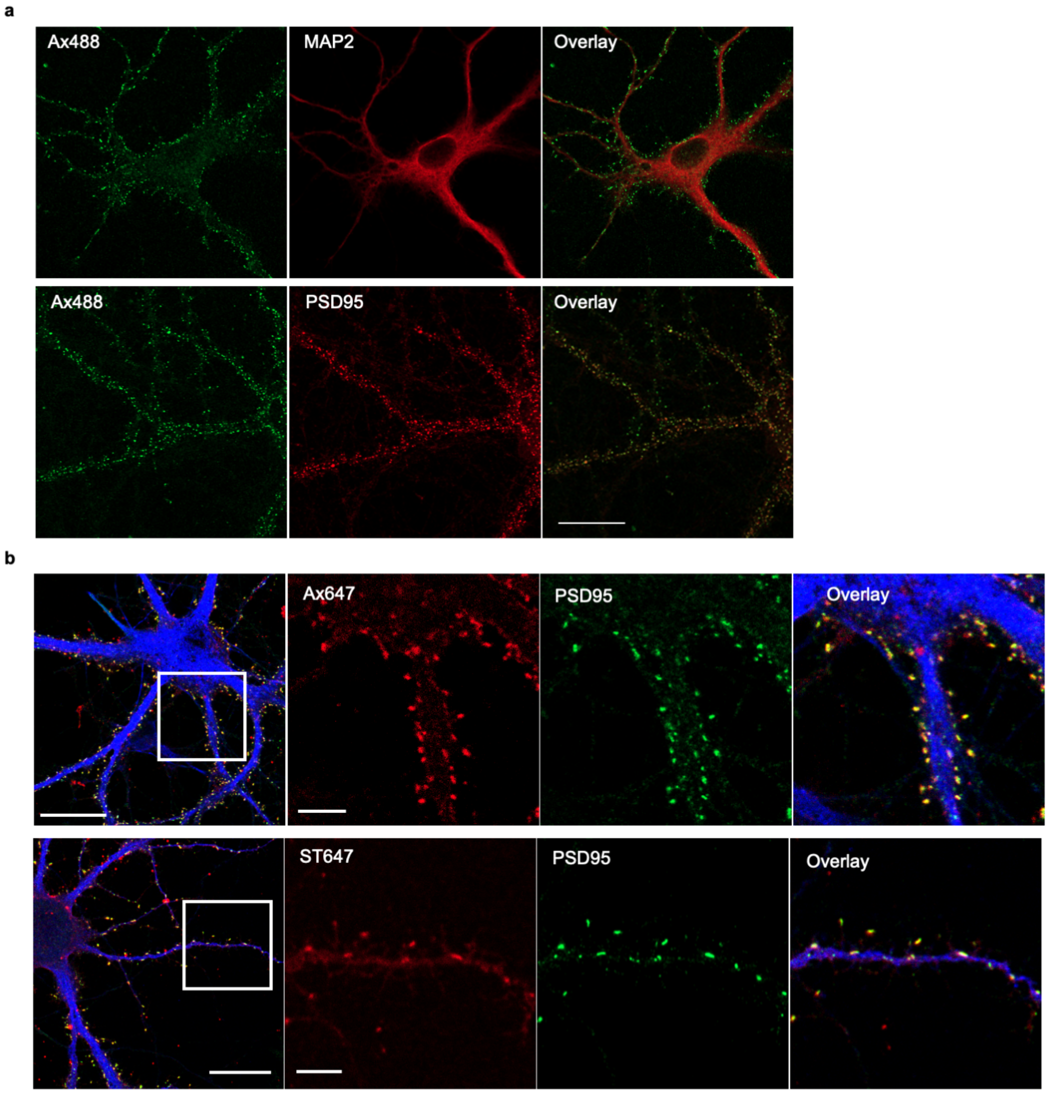


**Supplementary Figure 7 | Visualization of cell-surface AMPARs endogenously expressed in neurons by the two-step labeling.** (**a**) Whole images of immunostaining of cortical neurons after the two-step labeling. See also Figure 4c for expanded image. Labeling was conducted as described in Figure 4c. The neurons were fixed, permeabilized and immunostained using anti-MAP2 (upper) or anti-PSD95 antibody (lower). Scale bars, 20 μm. (**b**, **c**) Immunostaining of cortical neurons after the two-step labeling using **Tz(Ax647)** or **Tz(ST647)**. Labeling was conducted as described in Figure 4c. The neurons were fixed, permeabilized and immunostained using anti-PSD95 antibody. Scale bars, 20 μm or 5 μm in whole or expanded images, respectively.

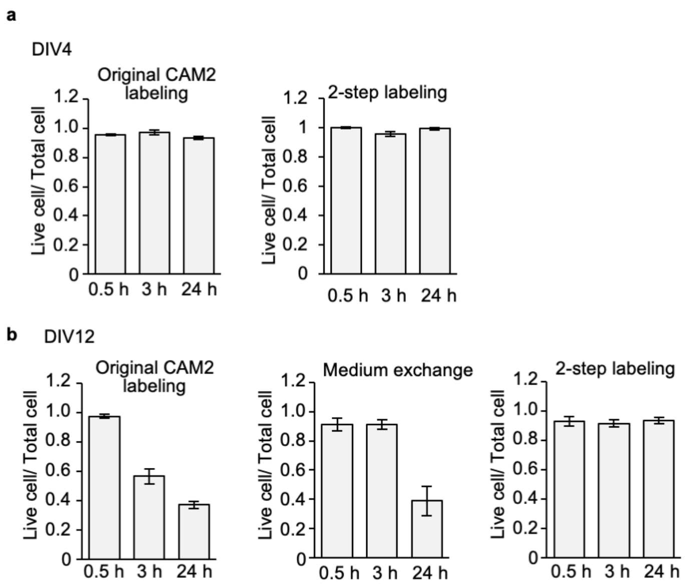


**Supplementary Figure 8 | Influence of two-step labeling process on the viability of cultured neurons.** (**a**, **b**) Effects of original CAM2 labeling process, medium exchange procedure into serum-free medium, two-step labeling process for immature cortical neurons (DIV4) in (**a**) or for mature cortical neurons (DIV12) in (**b**). Viability of neurons were evaluated by the Calcein AM Cell Viability Assay. In immature neurons, viability of neurons was not affected by these processes. In mature neurons, viability of neurons was not affected by the two-step labeling process, even though live cells decreased after 3 h of original CAM2 labeling process or after 24 h of medium exchange procedure. Raw fluorescent images are shown in Supplementary Figure 9.

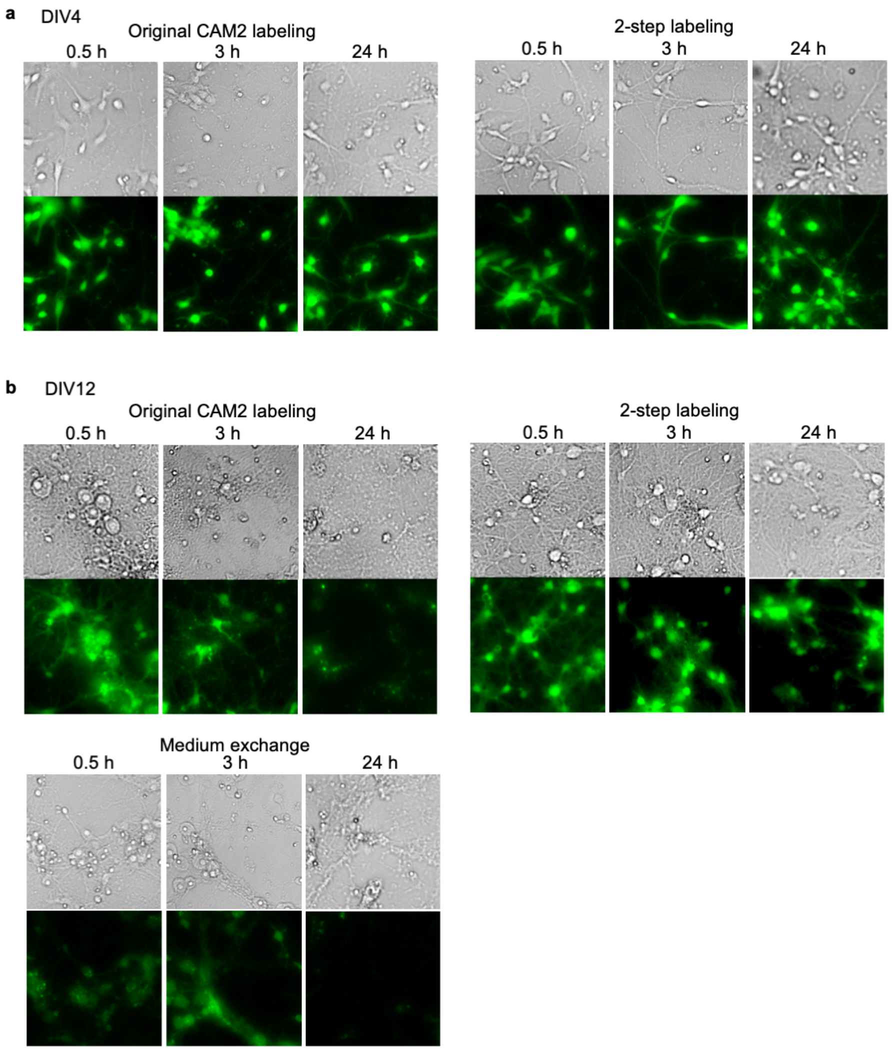


**Supplementary Figure 9 | Raw images of Calcein AM Cell Viability Assay.** (**a**, **b**) Epi-fluorescence images after Calcein AM staining are shown for immature cortical neurons (DIV4) in (**a**) or for mature cortical neurons (DIV12) in (**b**).

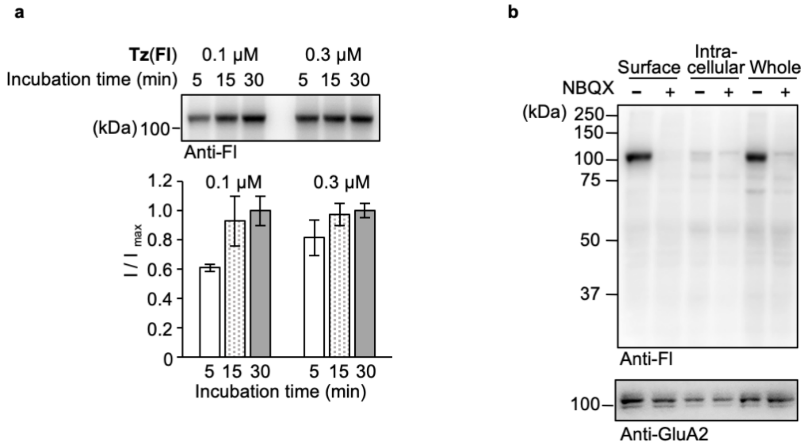


**Supplementary Figure 10 | Reaction kinetics of tetrazine ligation in cell lysates and a whole blot for surface, intracellular or whole-cell labeling in cultured cortical neurons.** (**a**) Reaction kinetics of tetrazine ligation in cell lysate evaluated by western blotting. After cell lysis of **CAM2(TCO)**-labeled neurons, each concentration of **Tz(Fl)** was added for 5, 15 or 30 min. Then, 10 μM TCO-PEG4-COOH was added for quenching **Tz(Fl)**. This result indicates that the tetrazine ligation was saturated within 15 min. (**b**) Whole blot for surface, intracellular or whole-cell. The sample was prepared as described in Figure 3g. Selective band corresponding to AMPAR was observed, which indicates high selectivity of the tetrazine ligation even in cell lysates. Data are represented as mean ± s.e.m.

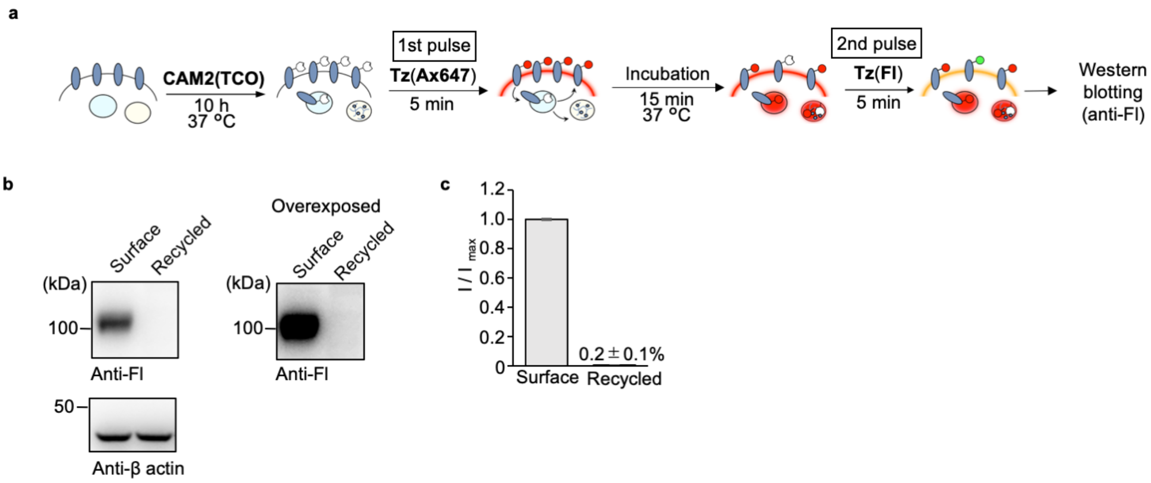


**Supplementary Figure 11 | Analyses of recycled AMPARs by pulse-chase-type analyses using the two-step labeling in HEK293T cells.** (**a**) Schematic illustration of the procedure is shown. (**b, c**) Analyses of recycled AMPARs by western blotting. In **b**, representative results of western blotting are shown. β-actin was utilized as the loading control. In the right, overexposed image of the western blotting by anti-Fl antibody is shown. Recycled AMPARs were not detected even in this exposed image. In **c**, Recycled AMPARs were quantified, which were normalized by that for surface labeling (n = 3). This clearly indicates that recycling of AMPARs were not observed in HEK293T cells. Data are represented as mean ± s.e.m.

**Supplementary Methods**

**Confocal live cell imaging of AMPARs labeled by CAM2(Ax488)**

HEK293T cells were co-transfected with GluA2 $^{flip}$(Q) and mCherry-F as a transfection marker. GluA2-expressing HEK293T cells were treated with 2 μM **CAM2(Ax488)** in serum free DMEM-GlutaMAX at 17 °C or DMEM-GlutaMAX supplemented with 10% dialyzed FBS at 37 °C for 4 h. The cells were washed 3 times with HBS. Confocal live imaging was performed with a confocal microscope.

**Fluorescent spectra measurements.**

Fluorescence spectra were measured with a Shimadzu RF-6000 fluorescence spectrometer using a quartz cell with 0.2 × 1.0 cm path length at excitation wavelengths of 430 nm (for Alexa 488) and 610 nm (for Alexa 647 and SeTau-647) at room temperature. Tz probes were dissolved in PBS at a concentration of 100 nM and their fluorescence was measured (TCO(-)). Subsequently, a 10-fold excess of TCO-PEG4-COOH was added into the quartz cell and incubated for 10 min at room temperature, after which the fluorescence spectrum was measured (TCO(+)).

**Confocal live imaging of AMPARs using various tetrazine probes in HEK293T cells.**

HEK293T cells were co-transfected with GluA2 and mCherry-F (for **Tz(Fl)** (Jena Bioscience)) or EGFP-F (for **Tz(Cy5)** (Jena Bioscience), **Tz(Ax647)**, **Tz(ST647)** and **Tz(Bt)** (Jena Bioscience)) as a transfection marker. GluA2-expressing HEK293T cells were treated with 2 μM **CAM2(TCO)** in the culture medium at 37 °C for 4 h. After removal of the culture medium, 100 nM **Tz(Fl)**, **Tz(Cy5)**, **Tz(Ax647)**, **Tz(ST647)** or **Tz(Bt)** was treated for 5 min in HBS at room temperature. In the case of the **Tz(Bt),** 1 μg/mL streptavidin-HiLyte647 (**SAv(Hilyte647)** (ANASPEC)) in HBS was treated at

37 °C for 10 min. The cells were washed 3 times with HBS, and then confocal live imaging was performed with a confocal microscope.

**Comparison of photostability between Ax647 and ST647 labeled on AMPARs.**

For photostability study of Ax647 and ST647 labeled on AMPARs, HEK293T cells were co-transfected with GluA2 and EGFP-F as a transfection marker. GluA2-expressing HEK293T cells were treated with 2 μM **CAM2(TCO)** in the culture medium at 37 °C for 4 h. After removal of the culture medium, 100 nM **Tz(Ax647)** or **Tz(ST647)** was treated for 5 min in HBS at room temperature and washed 3 times with HBS. Confocal live imaging was performed with a confocal microscope. Fluorescence images were acquired by excitation at 640 nm for Alexa 647 and SeTau-647 derived from diode lasers (laser power: 20.0%).

**Reaction kinetics of Tz ligation in cell lysate.**

**CAM2(TCO)**-labeled HEK293T cells or cortical neurons were lysed with RIPA buffer containing 1% protease inhibitor cocktail for 30 min at 4 °C, then the lysate was reacted with 0.1, 0.3 μM **Tz(Fl)** for 5, 15, 30 min at room temperature. To quench excess **Tz(Fl)**, 10 μM TCO-PEG4-COOH was added. For western blotting analysis, after chemical labeling, cells were mixed with 5× Laemmli sample buffer containing 250 mM DTT. The samples were applied to SDS-PAGE and electrotransferred onto PVDF membranes, followed by blocking with 5% nonfat dry milk in TBS containing 0.05% Tween 20. The Fl-labeled GluA2 was detected by chemiluminescence analysis using rabbit anti-fluorescein antibody (abcam, ab19491, 1:3,000) and anti-rabbit IgG-HRP conjugate (CST, 7074S, 1:3,000). The immunodetection of GluA2 was performed with a rabbit anti-GluA2/3 antibody (Millipore, 07-598, 1:3,000) and anti-rabbit IgG-HRP conjugate (CST, 7076S, 1:3,000). The signal was generated with ECL Prime (GE Healthcare) and detected with Fusion Solo S imaging system (Vilber Lourmat).

**Cell viability assay of matured and unmatured cultured cortical neurons.**

Primary cultures of cortical neurons were prepared as described above and used at 4 or 12 DIV. To label endogenous AMPARs by two-step labeling method, 12 μM **CAM2(TCO)** in 100 μL culture medium was gently added to the cortical neurons cultured in 500 μL medium on 24-well plates to a final concentration of 2 μM **CAM2(TCO)**. The cells were incubated for 10 h at 37 °C and further incubated for 0, 2 h and 24 h at 37 °C. For the second step labeling, the culture medium was removed and the cells were treated with 1 μM **Tz(Fl)** for 5 min in Neurobasal Plus medium at 37 °C. To quench excess **Tz(Fl)**, 1 μM TCO-PEG4-COOH in Neurobasal Plus medium was added.

To label endogenous AMPARs by original method[S1], 2 μM **CAM2(Ax488)** in culture medium with 10 mM HEPES and without B27 Plus supplement was added to the cortical neurons cultured on 24-well plates. The cells were incubated for 4 h at 17 °C, washed 3 times with HBS and further incubated for 0, 2 h and 24 h at 37 °C.

For cell viability assay, labeled cells were washed with HBS and incubated with 2 μM Calcein AM (Dojindo) and 50 μg/mL Hoechst33342 (Dojindo) in HBS for 15 min at 37 °C. Fluorescence live imaging was performed with a fluorescent microscope (IX71, Olympus) equipped with a 20×, numerical aperture (NA) = 1.4 objective.

The cell viability rate was calculated according to the following formula:

Live cell / Total cell = (Number of Calcein positive cells) / (Number of Hoechst33342 positive cells)

**Quantification of recycled AMPARs in HEK293T cells.**

GluA2-expressing HEK293T cells were labeled with 2 μM **CAM2(TCO)** in culture medium in 37 °C for 4 h. For the second step, the culture medium was removed and the cells were treated with 1 μM **Tz(Ax647)** for 5 min in the culture medium at 37 °C. To quench excess **Tz(Ax647)**, 1 μM TCO-PEG4-COOH in the culture medium was added.

After incubation at 37 °C for 15 min, recycled AMPARs were labeled with 1 μM **Tz(Fl)** for 5 min in PBS. To quench excess **Tz(Fl)**, 1 μM TCO-PEG4-COOH in PBS was added. Cell lysis and western blotting were performed as described in "Reaction kinetics of Tz ligation in cell lysate". The target bands were manually selected, and the intensity were calculated with ImageJ software, background intensity was manually subtracted by selecting a region with no bands around the target bands.

### Synthesis and Characterization

#### General materials and methods for organic synthesis

All chemical reagents and solvents were purchased from commercial sources (FUJIFILM Wako pure chemical, TCI chemical, Sigma-Aldrich, Sasaki Chemical) and were used without further purification. Thin layer chromatography (TLC) was performed on silica gel 60 F254 precoated aluminum sheets (Merck). Chromatographic purification was performed using flash column chromatography on silica gel 60 N (neutral, 40–50 μm, Kanto Chemical). $^{1}$H-NMR spectra were recorded in deuterated solvents on a Varian Mercury 400 (400 MHz) or JEOL JNM-ECA (600 MHz). Chemical shifts were referenced to residual solvent peaks or tetramethylsilane ($\delta$ = 0 ppm). Multiplicities are abbreviated as follows: s = singlet, d = doublet, t = triplet, m = multiplet, brs = broad singlet. High resolution mass spectra were measured on an Exactive (Thermo Scientific) equipped with electron spray ionization (ESI). Reversed-phase HPLC (RP-HPLC) was carried out on a Hitachi Chromaster system equipped with a diode array, and an YMC-Pack Triart C18 or ODS-A column.

### Synthesis of CAM2(TCO)

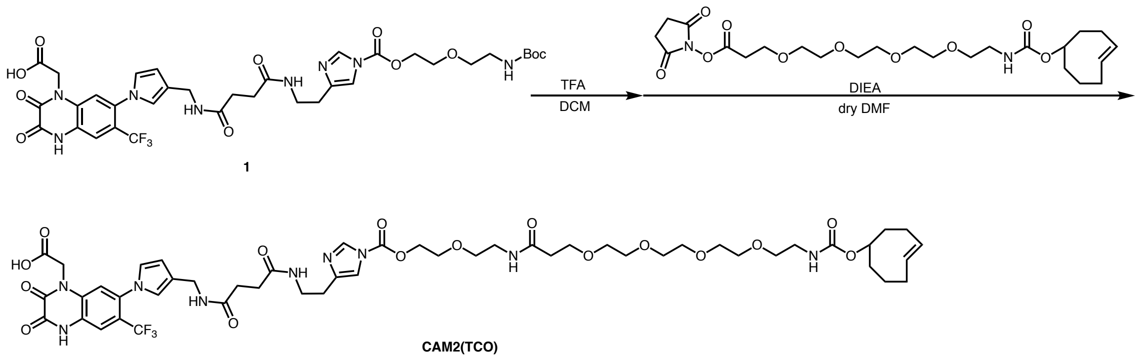


A solution of **1**[S1] (6.0 mg, 7.4 μmol) and TFA (0.5 mL) in dry DCM (0.5 mL) was stirred at room temperature for 4 h under $N_2$ atmosphere. After removal of the solvent by evaporation, the residual TFA was azeotropically removed with toluene (×3). The crude was used for the next step without further purification. A solution of the crude, TCO-PEG4-NHS (5.0 mg, 9.7 μmol) and DIEA (10 μL, 57 μmol) in dry DMF (0.5 mL) was stirred at room temperature for 13 h under $N_2$ atmosphere. The reaction mixture was purified by RP-HPLC (ODS-A, 250 x 25 mm, mobile phase; $CH_3CN$ : 10 mM $AcONH_4$ aq. = 10:90 for 5 min to 50:50 until 60 min (linear gradient over 55 min), flow rate; 10 mL/min, detection; UV (220 nm)), giving **CAM2(TCO)** (2.4 mg, 3.9 μmol, 53% yield in 2 steps) as a transparent oil. HR-ESI MS m/z calcd for $[M+H]^+$ 1106.4652, found 1106.4627.

### Synthesis of Tz(Ax647)

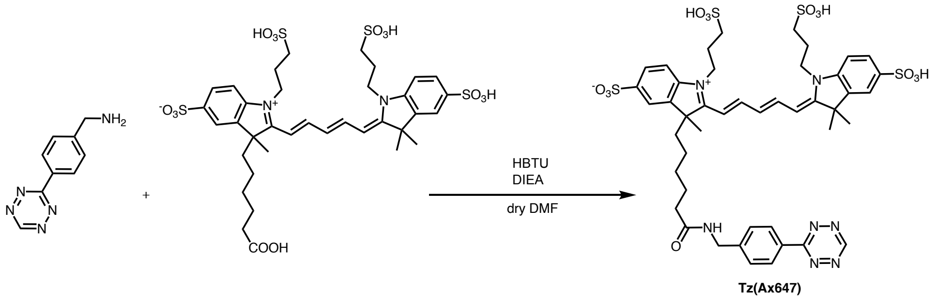


A solution of Alexa647 Carboxylic Acid (2.0 mg, 1.8 μmol), tetrazine benzylamine (1.2 mg, 5.3 μmol), HBTU (1.2 mg, 3.1 μmol) and DIEA (2.4 μL, 14 μmol) in dry DMF

(0.4 mL) was stirred at room temperature for 10 h under $N_2$ atmosphere. The reaction mixture was purified by RP-HPLC (ODS-A, 250 x 25 mm, mobile phase; $CH_3CN$ : 10 mM AcONH4 aq. = 5:95 for 5 min to 50:50 until 60 min (linear gradient over 55 min) , flow rate; 10 mL/min, detection; UV (220 nm)), giving **Tz(Ax647)** (0.8 mg, 0.15 μmol, 28%) as a blue solid. HR-ESI MS m/z calcd for $[M]^+$ 1028.2657, found 1028.2659.

**Synthesis of Tz(ST647)**

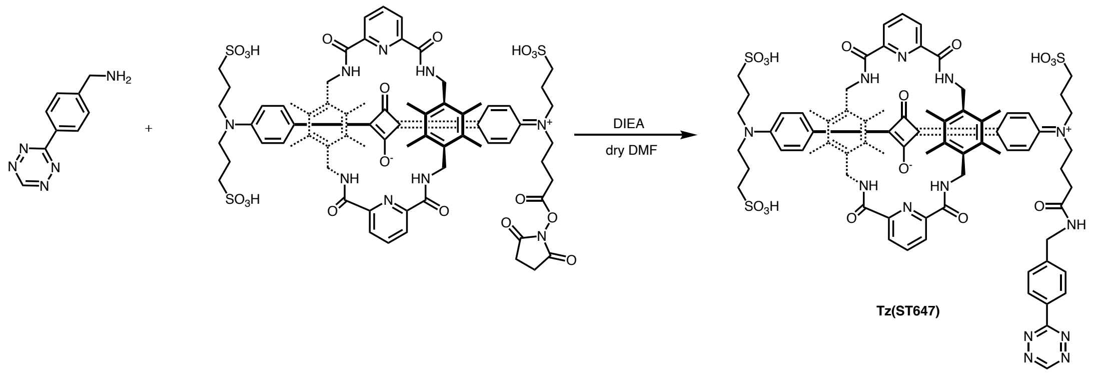


A solution of SeTau-647-NHS (1.0 mg, 0.54 μmol), tetrazine benzylamine (0.36 mg, 1.62 μmol) and DIEA (10 μL, 57 μmol) in dry DMF (1 mL) was stirred at room temperature for 3 h under $N_2$ atmosphere. The reaction mixture was purified by RP-HPLC (ODS-A, 250 x 10 mm, mobile phase; $CH_3CN$ : 10 mM $AcONH_4$ aq. = 5:95 to 50:50 (linear gradient over 60 min) , flow rate; 3.0 mL/min, detection; UV (220 nm)), giving **Tz(ST647)** (0.3 mg, 0.15 μmol, 28%) as a dark green solid HR-ESI MS m/z calcd for $[M-3H]^{3-}$ 509.5060, found 509.5061.

**Synthesis of CNM(TCO)**

Synthesis of compound **4**

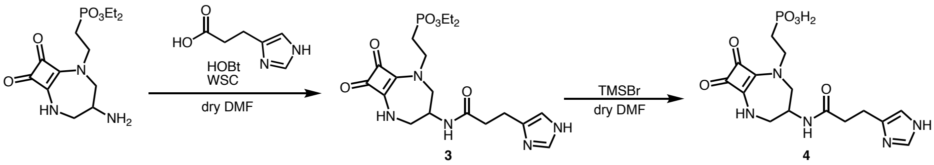


A solution of deamino-histidine (31 mg, 0.23 mmol), compound **2**[S2] (75 mg, 0.23 mmol), WSC・HCl (52 mg, 0.27 mmol) and HOBt (42 mg, 0.27 mmol) in dry DMF (4

mL) was stirred for 20 h. The crude was purified by flash column chromatography ($CHCl_3$ : MeOH = 2 : 1 + 1% $NH_3$ solution), giving compound **3** as a white solid (65 mg, 0.14 mmol, 63%). A solution of compound **3** (60 mg, 0.13 mmol) and TMSBr (121 µL, 0.92 mmol) was stirred in dry DMF (5 mL) at 60 °C for 7 h. After removal of the solvent by evaporation, the reaction mixture was purified by RP-HPLC (column; YMC-pack ODS-A, 250 ×10 mm, mobile phase; $CH_3CN$ (0.1% TFA): $H_2O$ (0.1% TFA) = 0:100 to 15 :85 (linear gradient over 30 min), flow 3.0 mL/min, detection; UV (220 nm)), giving **4** as a white solid (6.6 mg, 15.9 µmol). $^1$H-NMR (400 MHz, $CD_3OD$) $\delta$ 8.62 (s, 1H), 7.28 (s, 1H), 4.29-4.17 (m, 2H), 3.83-3.65 (m, 4H), 4.32 (d, $J$ = 13.2 Hz), 3.01 (t, $J$ = 7.2 Hz, 2H), 2.70 (d, $J$ = 7.2 Hz, 1H), 2.07-1.97 (m, 2H). HR-ESI MS m/z calcd for $[M+H]^+$ 398.1224, found: 398.1223.

**Synthesis of compound 6**

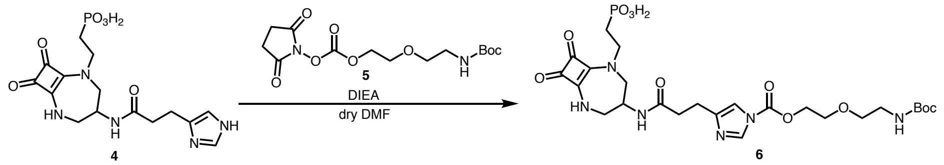


A solution of **4** (10.2 mg, 18.5 µmol), **5**[S1] (16 mg, 46.3 µmol) and DIEA (12.8 µL, 74 µmol) in dry DMF (1.6 mL) was stirred at room temperature for 5 h under $N_2$ atmosphere. The reaction mixture was purified by RP-HPLC (column; YMC-pack ODS-A, 250 ×10 mm, mobile phase; $CH_3CN$: 10 mM $AcONH_4$ aq. = 0 : 100 to 40 : 60 (linear gradient over 40 min), flow; 3.0 mL/min, detection; UV (220 nm)), followed by lyophilization to give **6** as a white solid (8.9 mg, 14.1 µmol). $^1$H-NMR (400 MHz, $CD_3OD$) $\delta$ 8.17 (s, 1H), 7.35 (s, 1H), 4.56 (br s, 2H), 4.28 (s, 1H), 4.05-3.74 (m, 7H), 3.56 (t, $J$ = 5.6 Hz, 2H), 3.40 (d, $J$ = 13.6 Hz, 1H), 3.23 (t, $J$ = 5.6 Hz, 2H), 3.56 (t, $J$ = 5.6 Hz, 2H), 2.62 (t, $J$ = 7.2 Hz, 2H), 2.00-1.92 (m, 2H), 1.42 (s, 9H) HR-ESI MS m/z calcd for $[M+H]^+$ 629.2331, found: 629.2330.

**Synthesis of CNM(TCO)**

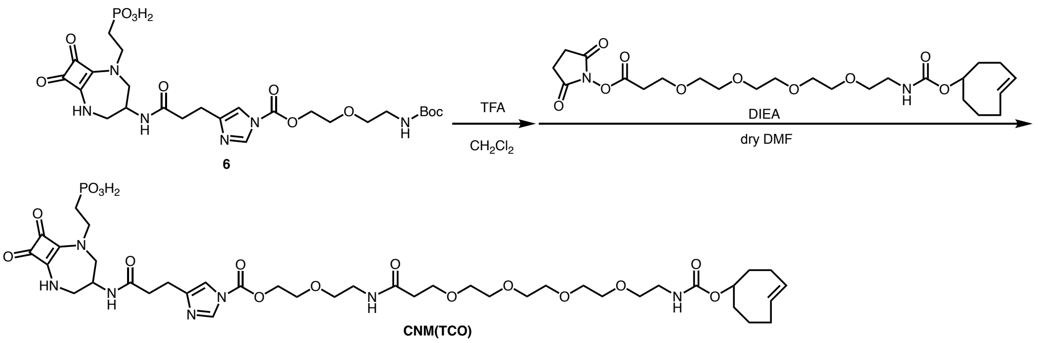


A solution of **6** (8.9 mg, 14.1 μmol) in $CH_2Cl_2$ (2 mL) and TFA (1 mL) was stirred at room temperature for 2.5 h under $N_2$ atmosphere. After removal of the solvent by evaporation, the residual TFA was azeotropically removed with toluene (×3). The crude was used for the next step without further purification. To a solution of the crude and DIEA (26 μL, 141 μmol) in dry DMF (1.2 mL) was added TCO-PEG4-NHS (10 mg, 19.4 μmol) in dry DMF (0.4 mL) and the mixture was stirred at room temperature for 5 h under $N_2$ atmosphere. Purification by RP-HPLC (column; YMC-pack ODS-A, 250 ×25 mm, mobile phase; $CH_3CN$: 10 mM $AcONH_4$ aq. = 0:100 to 50:50 (linear gradient over 50 min), flow 10.0 mL/min, detection; UV (220 nm)) followed by lyophilization gave **CNM(TCO)** as a white solid (3.0 mg, 3.2 μmol, 22% in 2 steps). HR-ESI MS m/z calcd for $[M+H]^+$ 928.4063, found: 928.4063.
